# CsHscB as a novel TLR2 agonist from carcinogenic liver fluke *Clonorchis sinensis* modulates host immune response

**DOI:** 10.1101/858670

**Authors:** Chao Yan, Fang Fan, Yu-Zhao Zhang, Jing Wu, Xin Dong, Hai-Liang Liu, Chun-Yang Fan, Qian Yu, Liang Wang, Xiang-Yang Li, Yu-Gang Wang, Jia-Xu Chen, Ren-Xian Tang, Kui-Yang Zheng

## Abstract

Clonorchis sinensis-a fluke dwelling on the intrahepatic bile ducts causes clonorchiasis. During C. sinensis infection, worm-host interaction results in activation of PRRs and further triggers immune responses which determine the outcome of infection. However, the mechanisms by which pathogen-associated molecules patterns from C. sinensis interacted with TLRs were poorly understood. In the present study, we identified a ∼34 kDa lipoprotein CsHscB from C. sinensis which physically bound with TLR2. We also found that recombinant CsHscB (rCsHscB) potently activated macrophage to express various proteins including TLR2, CD80, MHCII, and cytokines like IL-6, TNF-α, and IL-10 in a TLR2-dependent manner but rCsHscB failed to induce IL-10 in macrophages from *Tlr2*^-/-^ mice. Moreover, ERK1/2 activation was required for rCsHscB-induced IL-10 production in macrophages. In vivo study revealed that rCsHscB triggered a high induction of IL-10 in the wild-type (WT) but not in *Tlr2*^-/-^ mice. Our data thus demonstrate that rCsHscB from C. sinensis is an unidentified TLR2 agonist with immune regulatory activities, and may have some therapeutic implications in future beyond parasitology.

## Introduction

During helminth infection, the complex host-parasite interaction triggers host immune responses which ultimately drive the resistance to infection or immune evades accompanying with the course of immunopathogenesis. For this view, type 2 immune responses including IL-4, IL-9, IL-5 and IL-13 secreted by ILC2, Th2 or alternatively activated macrophage (AAM or M2) are typically considered as protective immunity against helminths to results in parasite expulsion ultimately [1]. However, the regulatory cells (Treg, Breg, ILCreg, M2c etc) can produce the regulatory cytokines (IL-10, etc) to ameliorate the bias of type II immune responses, which appears to be mainly responsible for the worms survival with the limited immunological damages and further establishment of chronic infection [2]. Further studies have demonstrated that MAPK (such as ERK, p38) and NF-κB signaling (NF-κB p50 homodimers) contribute to the mechanisms that control the production of IL-10 [3, 4]. However, the mechanisms by which the complex immune responses are initiated and finely-orchestrated remains poorly elucidated.

Toll-like receptors represent one of most important pattern recognition receptors (PRRs) that sense the conserved pathogen products (also called pathogen-associated molecular pattern, PAMP) from worms or alarming (also called dangers-associated molecular pattern, DAMP) sourced from damage tissues in the early event of infection. For example, TLR2 collaborated with TLR1 or TLR6 recognize triacylated or diacylated lipoproteins, respectively and thereby activate signal transduction cascades to result in the expression of pro-inflammatory or anti-inflammatory mediator genes [5-7]. So far several TLR2 ligands from *S. mansoni, Wolbachia*-endosymbiotic bacteria of *Brugia malayi* have been identified and demonstrated as potent immune regulators to determine the polarization of immune and even the outcome of helminth infection. For example, lysophosphatidylserine (Lyso-PS) from *S. mansoni* bound with TLR2 on dendritic cells allows DC to train IL-10 producing Tregs, which enables the long term survival of the parasite, as well as ameliorates of immunopathogenesis due to polarized type 2 immune responses [8]. Diacyl WoLP sourced from *Wolbachia* induces dendritic cell maturation and activation as well as drives CD4 T cell polarization and antibody switching in a TLR2-dependent manner [9].

Clonorchiasis caused by *Clonorchis sinensis* remains a major parasitic disease in eastern Asia such as China, Korea, Vietnam and eastern Rusia [10]. There are approximately 15 million people infected worldwide whereas 12.5 million people are distributed in China, posing a severe public health issue in these regions. The adult worms dwelling on the intrahepatic bile duct cause cholelithiasis, cholangitis, cholecystitis, biliary fibrosis and even cirrhosis due to its long-term survival. Additionally, chronic infection with this fluke has been shown to cause cholangiocarcinoma (CCA) and *C. sinensis* is now defined as Group 1 human biological agents (carcinogens) by International Agency of Research on Cancer (IARC) due to sufficient pieces of evidence in human [11, 12]. Previous studies have shown that the components of *C. sinensis* excretory/secretory products (ESPs) and crude antigen (CA) can potently induce a type 2 or a mix type1/type2 immune responses *in vitro* [13-15]. *In vivo* study, during *C. sinensis* infection, the interaction between worms and host immune cells also potently drives type I immune responses with type 2 becoming more prevalent after worms are well-developed in susceptible hosts [16]. Furthermore, our previous study also showed that the expression of TLR2 is dramatically changed with the prolonged infection, which suggested that TLR2 might be involved in this dramatically immunological changing [17]. However, the mechanisms that account for this phenotypic shifting is poorly understood so far. In view of this background, the objectives of the present study were to identify the molecules from *C. sinensis* that are responsible for activation of TLR2 and investigate its possible effects on the activation on macrophage. In our present study, we identified a lipoprotein-rCsHscB interacted with TLR2 acting as an unidentified TLR2 agonist induce the activation of macrophage secreting high levels of pro-inflammatory and anti-inflammatory cytokines in a TLR2-dependent manner. Our present study will contribute to a better understanding of the interaction between the *C. sinensis* and host cells. In addition, in view of regulatory immune capacities of rCsHscB, our study also provides an alternative therapeutic approach for implications beyond parasitology.

## Materials and Methods

### Ethics

Animal care and all experimental perform in this study were conformed to the guidelines of the National Laboratory Animal Center. The main procedures and protocol were reviewed and approved by the Animal Care and Use Committee of Xuzhou Medical University License (2016-SK-03).

### Mice

Male C57BL/6 mice (specific pathogen-free, SPF) aged 8 weeks (20-22g) were purchased from the Beijing Vital River Laboratory Animal Technology Co. Ltd. The mice were group-housed in a specific pathogen-free condition with temperature-control room (25 °C). All mice were given standard chow diet and tap water *ad libitum*.

To obtain *C. sinensis*-positive sera, BALB/c mice were orally infected by 45 nangyou and the mice were sacrificed on 28 days and 56 day post-infection (p.i.), the sera from *C. sinensis*-infected mice and no-infected mice were collected for further use.

Mice were immunized with around 10 μg of *C. sinensis* crude antigen in IFA. Two booster doses in IFA were injected in 15 days interval. Titers of antibody against to *C. sinensis* crude antigen were determined by ELISA.

### Preparation of rCsHscB and control protein

rCsHscB and control protein (only His-tagged protein encoded by pET-28a vector without CsHscB open reading frame) were routinely expressed by *E. coli* (Ec). rCsHscB was purified by nickel-affinity and ion-exchange chromatography. For more details, see the Supplementary Material.

### Development of specific rCsHscB polyclonal antibody

The Ab to the rCsHscB protein was generated in rabbits that were maintained in the animal house facility of Xuzhou Medical University. In brief, rabbits were immunized with around 10 μg of rCsHscB in IFA. Two booster doses in IFA were injected in 15 days interval. After measuring the rCsHscB-specific Ab titer by ELISA, animals were sacrificed at day 45 to collect and separate sera. The poly-antibody against rCsHscB was purified by metal affinity chromatography. In an immunoblot, the Ab raised against the rCsHscB protein specifically recognized single band of ∼36 kDa.

### Immunohistochemistry

rCsHscB was stained on paraffin-embedded adult worm *C. sinensis* by immunohistochemistry using the affinity-purified anti-rCsHscB antibody. Reactivity was detected using Dako REAL™ EnVision™ Detection System ((Dako, Glostrup, Denmark). Sections were counterstained with hematoxylin and photographed by a microscope.

### Cell culture and stimulation

Mouse mononuclear macrophage leukemia cells RAW264.7 with 5∼10 passages were cultured in DMEM (Hyclone, US) containing 10% fetal bovine serum (FBS) (Serana, AUS), 1% penicillin/streptomycin (Beyotine, China) in a humidified atmosphere with 5% CO_2_ at 37°C. RAW264.7 cells were stimulated by rCsHscB (5∼20 μg/ml) for 6 h, 12 h and 24 h. Supernatants were collected for assessing the concentrations of cytokines using ELISA. For TLR2 blocking assay, RAW 264.7 cells were pretreated with MAb-mTLR2 (2 µg/ml) or isotype (Invivogen, US) for 2 h. The cells were then stimulated by rCsHscB (20 μg/ml) or Pam_3_CSK_4_ (200 ng/ml) (Invivogen, US) for 24 h in a humidified atmosphere with 5% CO_2_ at 37 °C. The supernatants and cultured cells were collected for flow cytometry assays for ELISA, respectively.

Bone marrow cells were obtained from the long bones of 8- to 10-week-old C57BL/6 mice (WT or *Tlr2*^-/-^). Bone marrow cells were cultured in the presence of M-CSF (20 ng/mL) (PeproTech, USA) for six days to generate the bone marrow-derived macrophages (BMDMs). BMDMs were cultured in DMEM (Hyclone, US) containing 10% fetal bovine serum (FBS) (Serana, AUS), 1% penicillin/streptomycin (Beyotine, China) in a humidified atmosphere with 5% CO_2_ at 37°C. Thereafter, BMDMs were stimulated by rCsHscB (5∼40 μg/ml) or production of *E. coli* transfected by pET-28 control vectors for 24h and supernatants were obtained for determining the concentration of IL-10 using ELISA. For ERK1/2 inhibitor assay, PD98059 (1 µM) (Sigama, US) was pre-incubated with cells for 2 h, BMDMs from WT or Tlr2-/-mice were stimulated by rCsHscB (20 μg/ml) or Pam_3_CSK_4_ (200 ng/ml) (Invivogen, US) and supernatants were used for ELISA.

### Western blotting analysis

Total cell lysates or rCsHscB were separated by 10% SDS-polyacrylamide gel electrophoresis (PAGE) and transferred onto Immobilon-P Transfer Membranes (PVDF) (Millipore, USA). For detection specific antibodies against to rCsHscB in vivo, sera from C. sinensis-infected and non-infected mice, as well as C. sinensis crude antigens immunizing sera as primary antibodies for 12 h at 4°C, and then horseradish peroxidase-conjugated secondary antibody (Beyotine, China) were incubated. For detection of MAPK signaling, the PVDFs were blocked with 5% non-fat-milk in PBS-Tween (PBS-T) and incubated with anti-His (ZSGB-Bio, China), anti-phospho ERK (CST, US), anti-phospho p38 (CST, US), anti-TLR2 (CST, US), anti-β-actin (Beyotine, China), and horseradish peroxidase-conjugated secondary antibody (Beyotine, China). The PVDFs were visualized by ECL exposure to X-ray film. Densitometry analyses were performed by Image Lab software.

### Flow cytometry

Following stimulation, RAW264.7 were stained with TLR2 (eFlour 660), CD80 (PE), CD86 (eFlour 450), major histocompatibility complex class II (FITC), CD206 (PE/Cy7), CD11b (APC-Cy7). BMDMs were stained with TLR2 (eFlour 660), CD11b (APC-Cy7), F4/80 (Percp-Cy5.5). Antibodies were purchased from BD Pharmigen (US). Samples were analyzed with FlowJo software.

### ELISA

Supernatants from RAW264.7 or BMMs cultures were analyzed using commercially available ELISA kits for IL-10, IL-6 and TNF-α (all from eBioscience, San Diego, CA, US).

### Pull-down assay

The cells were stimulated by supernatant of lysate from *E. Coli* transfecting with Vector controls (pET-28, His-tagged control), pET-28a-CsHscB vectors (pET-28-CsHscB, unpurified), purified rCsHscB-stimulated cells, binding buffer and medium for 24 h, subsequently, the cells from each group were lysed for further use. The rCsHscB were incubated with Ni-NTA beads (QIAGEN, GER) for 12h at 4°C after the agaroses were balanced with binding buffer at 4 times in 4°C. rCsHscB immobilized on bead were incubated with total cell lysates (RAW264.7) for 12h at 4°C. The supernatant was discarded after centrifuged at 2500 rpm for 5 minutes in 4°C. The bead-bound proteins were subjected to 10% SDS-PAGE and then transferred electrophoretically to PVDF membranes. The membranes were incubated with anti-His antibody or anti-TLR2 antibody, followed by horseradish peroxidase-conjugated secondary antibody (Beyotine, China). The PVDFs were visualized by ECL exposure to X-ray film.

### Statistical analysis

All data were expressed as the mean ± standard error of the means (SEM). One-way ANOVA was used to analyze the significance of the differences between groups, followed by Tukey’s test using SPSS 13.0. For all tests, *P*<0.05 was considered statistically significant.

## Results and discussion

### Identification, characterization and immunogenicity of recombinant *C. sinensis* HscB

As most TLR2 agonists or ligands have been reported as lipoproteins or lipopeptides [18], to identify potential agonist of TLR2 sourced from *C. sinensis*, we collected all the amino acid sequences encoding *C. sinensis* proteins from the proteome data (http://www.ncbi.nlm.nih.gov/bioproject/PRJDA72781) and then putative lipoproteins from *C. sinensis* proteome were screened and predicted using a combination of DOLOP, lipoP and Lipo database as previously described [9]. We ultimately identified a lipoprotein named molecular chaperone HscB (CsHscB), which had 283 amino acids with three domains as followed: DnaJ, Co-chaperone HscB (COHscB) and C-terminal oligomerization (CTO) (Fig. 1A). Alignment of amino acid sequences analysis showed that the sequences of *C. sinensis* HscB had more than 90% similarities to *Opisthorchis viverrini* hypothetical protein (XP_009168973.1), but only had 40.91% similarities to the putative co-chaperone protein HscB from *Schistosoma mansoni* and 33.64% to the co-chaperone HscB from *Echinococcus granulosus* (Fig. 1B). The candidate lipoproteins were further selected for prediction of N-terminal signal peptide using SignaIP server 3.0 (http://www.cbs.dtu.dk/services/SignalP-3.0/) by a hidden Markov model (HMM) [19]. CsHscB had a signal peptide (the probability was 0.759, Fig. 1C) and the predicted cleavage sites was at between N-terminal 34 and 35 sites (Fig.1C).

**Figure 1.**
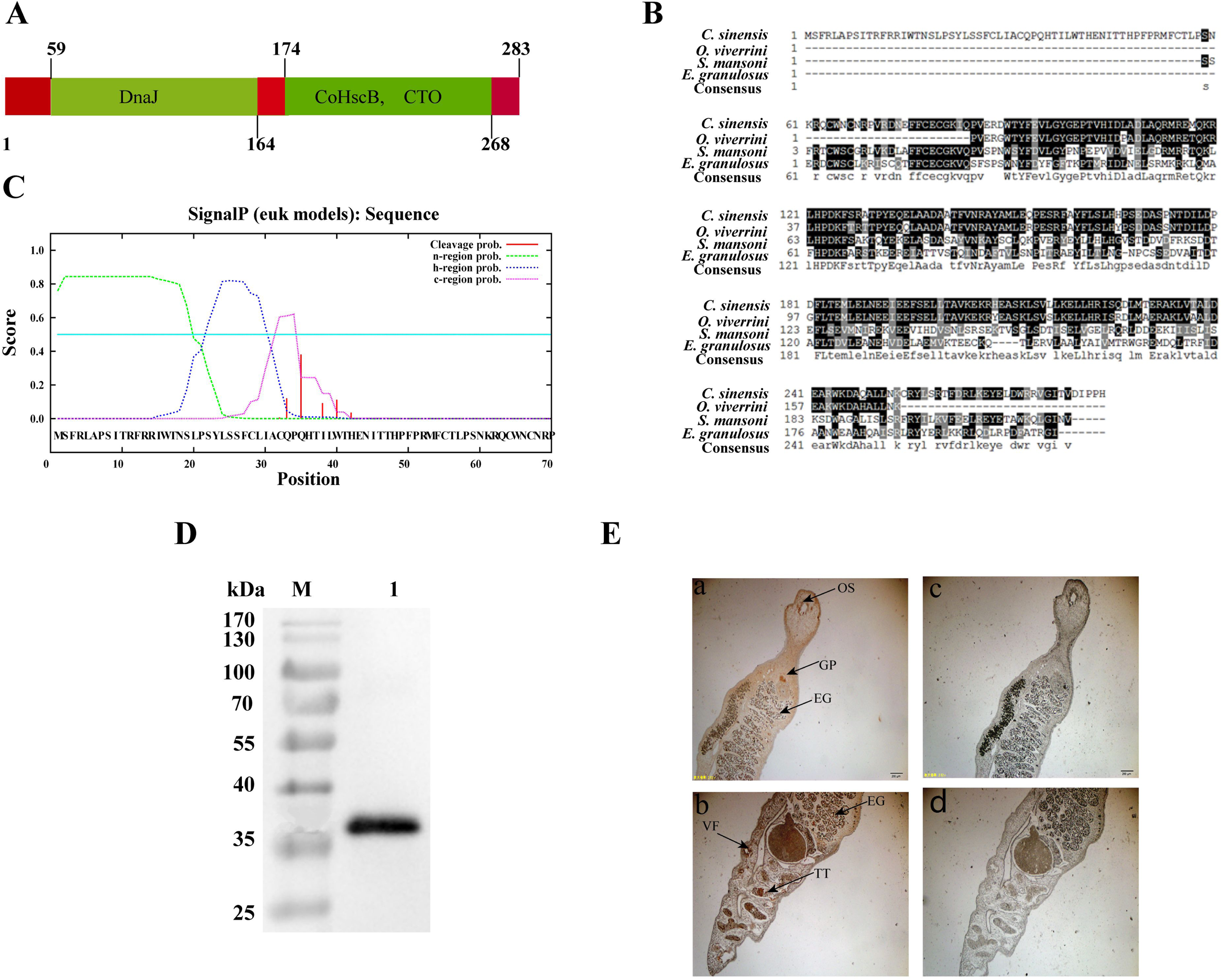
Identification, characterization and immunogenicity of *Clonorchis sinensis* HscB. **A** Secondary structure of CsHscB including three domains: DnaJ, Co-chaperone HscB (CoHscB) and C-terminal oligomerisation (CTO). **B** Multiple protein sequences alignment of CsHscB among different species of helminth. **C** Signal peptide identification of CsHscB using SignalP server 3.0. **D** Western-blot analysis of purified rCsHscB using anti-His antibody. **E** Expression and distribution of rCsHscB in the *C. sinensis* adult worms using IHC. a∼b: rCsHscB on the worm bodies were detected using primary antibody of rCsHscB; c∼d: rCsHscB on the worm bodies were detected using IgG isotype as primary antibody for controls. Arrows indicates the expression of rCsHscB mainly on the oral sucker (OS), vitelline follicles (VF), genital pore (GP), testis (TT) and eggs (EG). Scale bars, 20 μm.

However, it was very difficult to isolate and purify CsHscB directly from the worms due to lack of the sufficient background information as well as the low yield. We therefore used a recombinant CsHscB (rCsHscB) that was routinely expressed by *E. coli* (Ec). rCsHscB was purified by nickel-affinity and ion-exchange chromatography, and the purified rCsHscB was assessed by western-blot (Fig 1D). The molecular weight of rCsHscB including a 6×his tag was approximate 36 kDa (Fig 1D). Furthermore, we prepared the specific rCsHscB antibody to examine the expression and distribution of CsHscB in worm body using immunohistochemistry (IHC). IHC data showed that CsHscB mainly expressed on the oral sucker (OS), genital pore (GP), vitelline gland (VF), ovary (OV), testis (TT) and eggs (EG) (Fig. 1E). It could be detected by the sera from *C. sinensis*-infected mice as well as *C. sinensis* crude antigen-immunized mice, suggesting that rCsHscB was recognized by pool of antibodies induced by *C. sinensis* and crude antigen as well. It is also suggested that CsHscB naturally existing in *C. sinensis*-infection mice and worm’s crude antigens can trigger host immune responses (Fig. S 1).

### rCsHscB induces the activation of macrophage and cytokine production

To test whether rCsHscB has the capacity to induce the activation of innate immune cells or not, we used a macrophage cell line-RAW 264.7 that were stimulated by various concentrations of rCsHscB at different time-points. Firstly, we test toxicity of rCsHscB to macrophages, lactate dehydrogenase (LDH) test showed that up to 20 µg /ml of rCsHscB protein displayed no cellular toxicity against macrophages (the data is not shown). Furthermore, endotoxin (LPS) in the purified rCsHscB was removed by Endotoxin Erasol Solution (Tiandz, Beijing, China) in order to exclude any potential effects of LPS produced during preparation of rCsHscB. The concentration of endotoxin was detected by Limulus Amebocyte Lysate (LAL) and rCsHscB solution with less than 0.1 EU/ml of endotoxin should be further studied. For assessment of the activation of macrophages, we detected activation markers of macrophages upon stimulation using flow cytometry. The data showed that stimulation of macrophage with rCsHscB (20 μg/ml) for 24 h augmented the surface expression of activation markers such as TLR2, CD80, CD86, MHCII, CD206 and CD11b (Fig. 2A∼F). We also detected these cytokines with various concentrations (5∼20 μg/ml) of rCsHscB at different time courses, it was shown that macrophages stimulated by 5∼20 μg/ml rCsHscB for 12 h produced high levels of TNF-α (Fig. 2G). In addition, rCsHscB with the concentration of 5∼10 μg/ml but not 20 μg/ml for 24 h still induced a robust secretion of TNF-α produced by macrophage (∼3 times greater than DMEM-stimulated cells, Fig. 2G). The cells also produced high levels of IL-6 under the stimulation with 5∼20 μg/ml rCsHscB for 12 h or 24 h, compared with medium-stimulated cells (Fig. 2H, *P*<0.05). With regard to IL-10, cells stimulated with 5∼20 μg/ml rCsHscB for 12 h or 24 h could produce a robust increase of IL-10, of note, the secretion of IL-10 in macrophage stimulated by 20 μg/ml rCsHscB for 24 h was more than 10 times greater than that of DMEM-stimulated cells (Fig. 2I). We also tested the levels of IL-4 and IL-12 produced by rCsHscB-stimulated macrophage, but the data showed that the macrophage stimulated by rCsHscB didn’t increase the production of IL-4 and IL-12 (the data is not shown).

**Figure 2.**
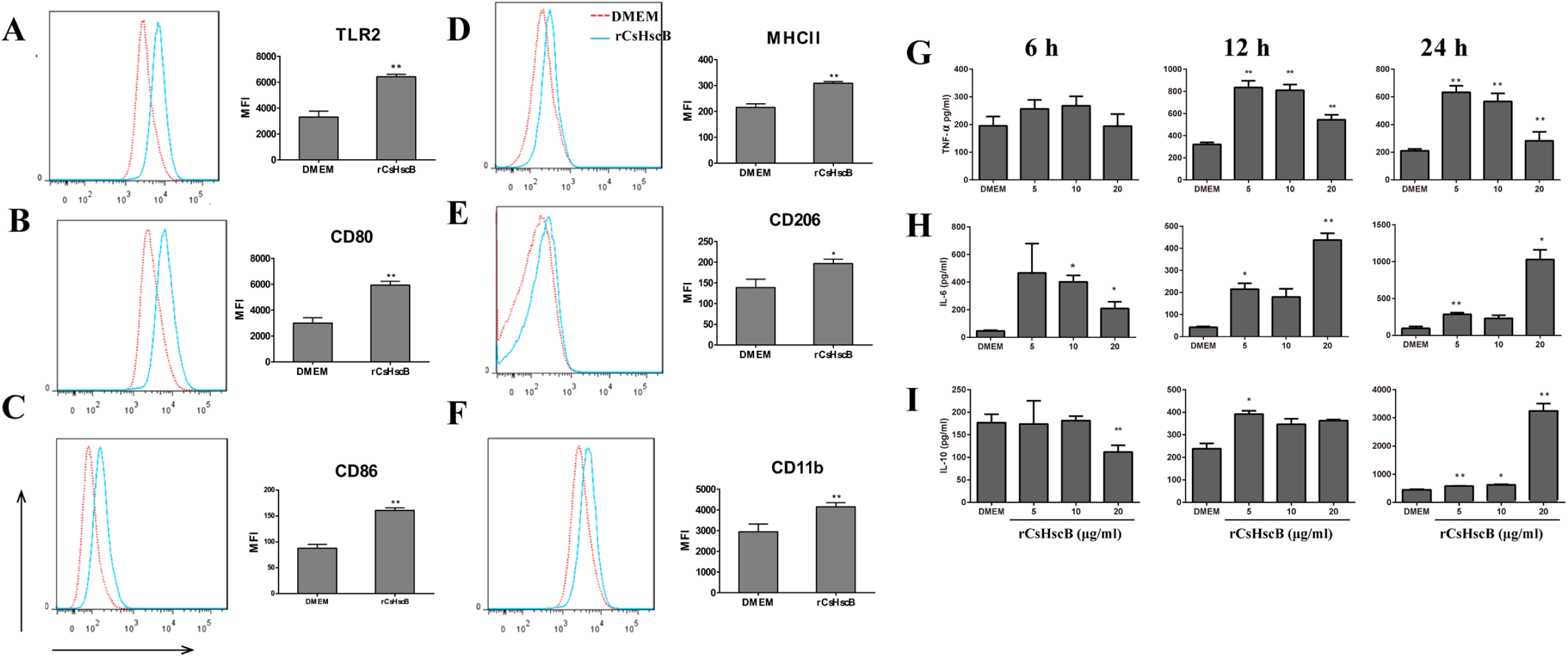
rCsHscB induces the activation of macrophage and cytokine production. **A**∼**F** rCsHscB increased the expression of macrophage activation markers by flow cytometry. **G∼I** productions of TNF-α(G), IL-6 (H) and IL-10 (I) were assayed for ELISA in the macrophage stimulated by indicated concentrations of rCsHscB for different time courses. Quantitative data are the mean±SE of three independent experiments, and all data shown are representative of at least three independent experiments. \**P* < 0.05, \*\**P* < 0.01, and \*\*\**P* < 0.001, stimulated cells *versus* those cultured in medium alone.

To exclude any potential effects of endotoxin and other potential component produced during preparation of rCsHscB on the activation of macrophage, we also compared the productions of *E. Coli* induced by pET-28a vector with or without CsHscB open reading frame (pET-28a-CsHscB or control vector), the production induced by control vector could not stimulate macrophage to secrete high levels of IL-10 and TNF-α (Fig. S2A and Fig. S2B). However, it seems that the cells that were stimulated by the production expressed by control vector also secreted a higher level of IL-6, compared with DMEM stimulated cells, although the level of IL-6 was still lower than that of pET-28a-CsHscB-induced cells, suggesting that the increased secretion of IL-6 may be not exclusively induced by rCsHscB (Fig. S2C). Together, these data demonstrate that rCsHscB induces the activation of macrophage and triggers a robust cytokines production by the macrophage.

### rCsHscB is an unidentified agonist for TLR2 to induce immune responses of macrophage

As the lipoproteins or lipopeptides have been reported as TLR2 agonist or ligand [18], we next test whether rCsHscB as an agonist for TLR2 to promote the activation of macrophage or not. Firstly, we performed *in silico* molecular docking using the crystal structure of the extracellular domain (ECD) of mouse TLR2 and modeled 3D structure of rCsHscB by homology modeling. Molecular docking showed that CsHscB could bind with TLR2 at its leucine-rich region (LRR) 11∼15 sites of ECD (Fig. 3A). To further ascertain whether rCsHscB physically interacts with the TLR2 molecule or not, we performed a pull-down assay using whole-cell extracts from RAW247.6 cells stimulated by rCsHscB. The cell extracts were incubated with rCsHscB immobilized on Ni-NTA beads. TLR2 Pull down assay revealed that supernatant of lysate from *E. Coli* transfecting with pET-28a-CsHscB vectors as well as purified rCsHscB proteins could pull-down TLR2 molecule as showed by western-blot (Fig. 3B line 2 and line 3), demonstrating that TLR2 and rCsHscB could be physically interacted. However, if rCsHscB (control vectors for example) absence, no bands were observed on the gel of western-blot, which suggested there is no interactions between TLR2 and other molecules except rCsHscB (Fig. 3B line 1 and line 4). Collectively, these data suggested that rCsHscB interacts specifically and predominantly with TLR2.

**Figure 3.**
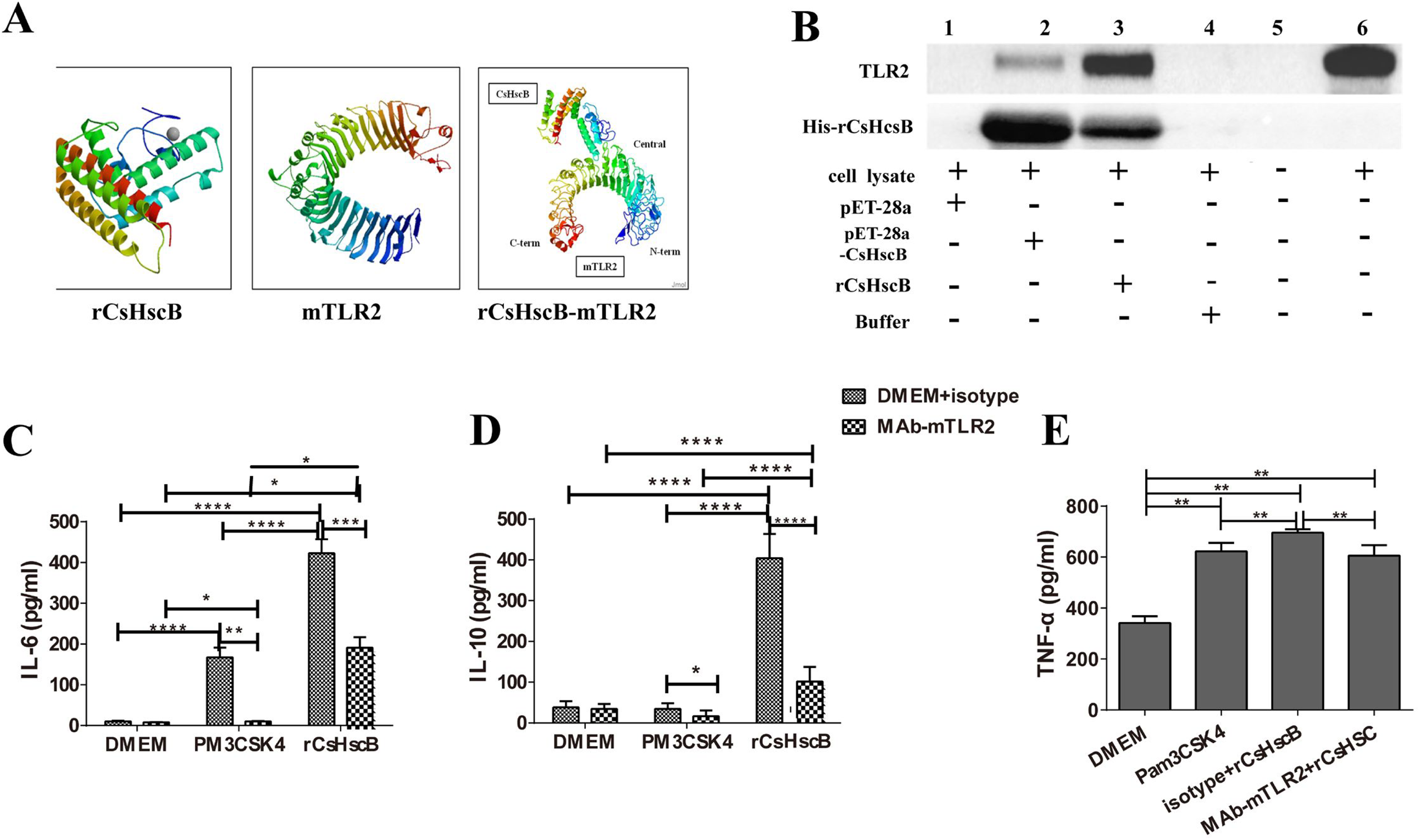
rCsHscB is a novel agonist for TLR2 to induce immune responses of macrophage. **A** Molecular docking analysis of binding TLR2 with CsHscB. **B** Pull-down assay analysis of interaction of TLR2 and rCsHscB. The cells stimulated by supernatant of lysate from *E. Coli* transfecting with Vector controls (pET-28, His-tagged control, Lane 1), pET-28a-CsHscB vectors (pET-28-CsHscB, unpurified, Lane 2), purified rCsHscB-stimulated cells (Lane 3), binding buffer (Lane 4) and medium (Lane 6) for 24 h, the cells were lysed and incubated with rCsHscB immobilized on Ni-NTA beads, and bead-bound proteins were loaded onto a gel for immunoblotting for TLR2 and rCsHscB, respectively. Lane 5 represents negative control for pull-down assay; **C∼E** The production of IL-6 (C), IL-10 (D) and TNF-α (E) were hindered in rCsHscB-stimulated cells when TLR2 was blocked by neutralizing antibody. PM_3_CSK_4_ were used as TLR2 ligand for positive controls. Quantitative data are representative of mean ± SEM of at least three independent experiments. \**P* < 0.05, \*\**P* < 0.01, and \*\*\**P* < 0.001 compared with indicated group.

To test whether rCsHscB induced the cytokines production is dependent on TLR2 or not, TLR2 was blocked by pretreating RAW 264.7 cells with anti-TLR2 antibody (T2.5) for 2 h prior to the addition of rCsHscB. The secretion of IL-6 and IL-10 induced by rCsHscB were almost abrogated following the addition of TLR2 blocking antibodies to the medium (Fig. 3D and Fig. 3E). Similarly, rCsHscB-induced TNF-α was also significantly abolished due to the presence of the TLR2 blocking antibody (Fig. 3F).

### rCsHscB-induced IL-10 production partly depends on phosphorylation of ERK1/2 but not p38 in RAW 264.7 cells

IL-10 has been known as one of the mechanisms that contribute to induced regulatory responses induced by helminth infection (30). As our and others’ previous studies suggest that IL-10 might play a regulatory role in immune responses during *C. sinensis* infection[17, 20-22], which highlight the significance of rCsHscB induced a strong IL-10 production in macrophage in our study, it is necessary to further study the mechanisms that IL-10 induced by rCsHscB. For this sense, to determine which downstream molecules mediated by TLR2 are responsible to robust rCsHscB-induced IL-10 production by macrophage, we screened the activation of transcription factors nuclear factor-κB (NF-κB), p38 mitogen-activated protein kinase and ERK1/2 in RAW264.7 cells using optimal concentrations (20 μg/ml) of rCsHscB during various time courses, western blot showed that rCsHscB induced a robust phosphorylation of ERK1/2 after 20∼30 min and then the levels of phosphorylation of ERK1/2 was attenuated during 60 min∼120 min following stimulation with rCsHscB (Fig. 4A). Surprisingly, there was no obviously activation of NF-κB nor p38 during these time courses. Furthermore, we also examined whether the rCsHscB-induced phosphorylation of ERK1/2 was mediated by TLR2 signaling. RAW264.7 cells were pretreated with blocking antibodies of TLR2 or with isotype control and phosphorylation of ERK1/2 was measured by western blot. Western blot showed that phosphorylation of ERK1 but not ERK2 was solely abolished following the addition of TLR2 blocking antibodies to the cultures, compared with isotype-matched control. Furthermore, we used a specific inhibitor for ERK1/2 (PD98059) to examine whether rCsHscB-induced cytokines was mediated by ERK signaling pathway or not. The RAW264.7 cells were pretreated with 10 µM PD98059 for 2 h, and then stimulated by 20 µg/ml rCsHscB for 24 h, the supernatants were collected for IL-10 detection. The data showed that the level of IL-10 was significantly decreased when ERK1/2 was inhibited by PD98059 in macrophage that was stimulated by rCsHscB for 24 h (Fig. 4C, *P*<0.001, ∼50% decreased).

**Figure 4.**
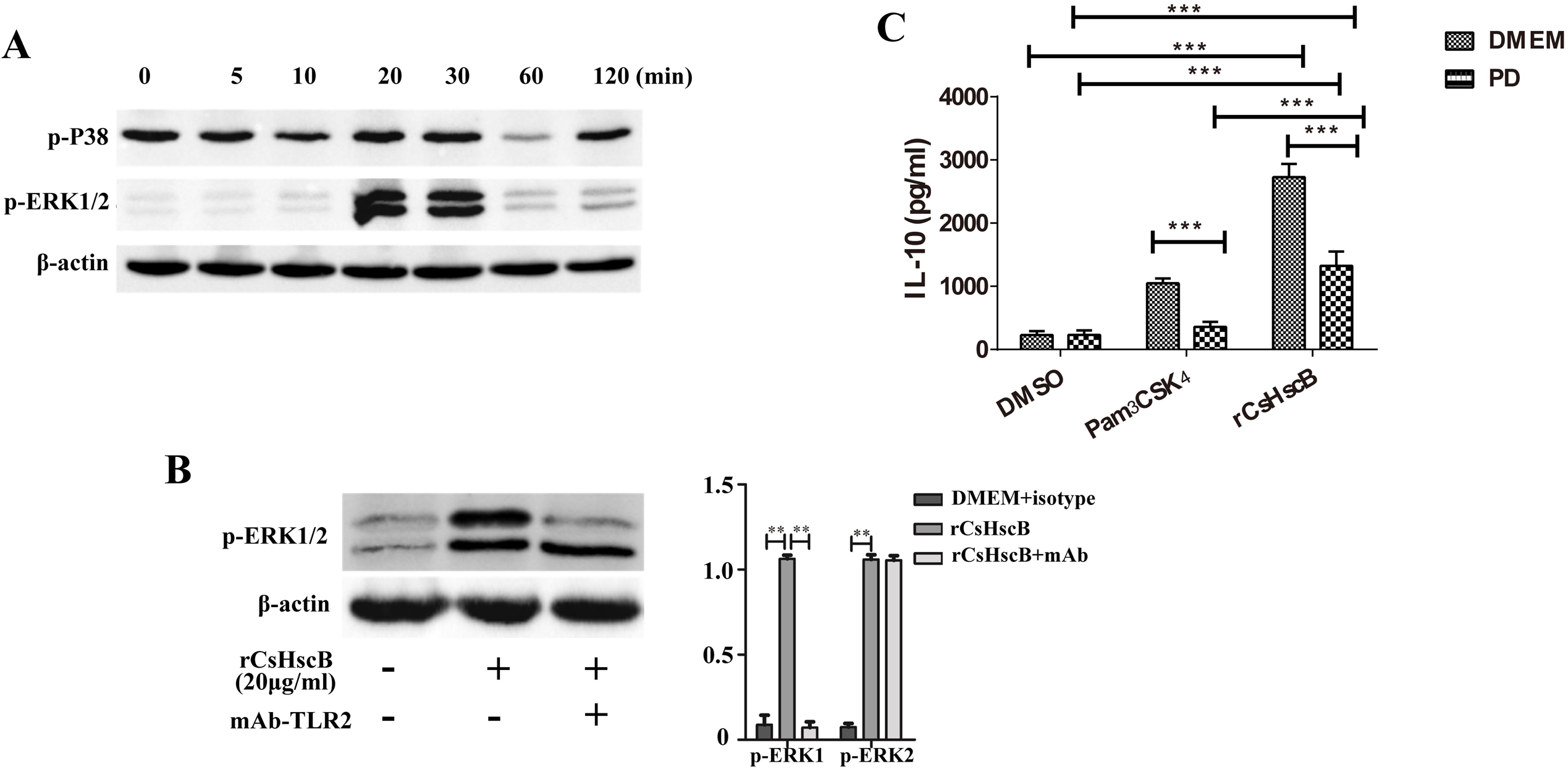
rCsHscB-induced IL-10 production partly depends on TLR2-meiated phosphorylation of ERK1/2 in RAW 264.7 cells. **A** ERK1/2 but not p38 MAPK was activated in rCsHscB-stimulated RAW 264.7 macrophage for 20∼120 min detected by western-blot. **B** phosphorylation of ERK1 but not ERK2 was attenuated in rCsHscB-stimulated cells when TLR2 was blocked by neutralizing antibody determined by western-blot. **C** ELISA analysis of IL-10 production in rCsHscB-stimulated RAW 264.7 cells the presence or absence of ERK1/2 inhibitor (PD98059). Quantitative data are representative of mean ± SEM of at least three independent experiments. \**P* < 0.05, \*\**P* < 0.01, and \*\*\**P* < 0.001 compared with indicated group.

### rCsHscB-induced IL-10 production depends on TLR2-mediated ERK1/2 signaling in bone marrow-derived macrophage

To ascertain the roles of TLR2-regulated ERK1/2 signaling in rCsHscB-induced IL-10 in macrophage, we induced bone marrow-derived macrophage (BMDM) from *Tlr2* wild type and *Tlr2*^-/-^ mice. Similar to our previous data, rCsHscB could potently induced a strong TLR2 expression on the surface of BMDM sourced from wild type mice (Fig. 5A, almost 2 fold changes) and the levels of IL-10 were significantly increased when BMDM cells from *Tlr2* wild-type mice were stimulated at various concentration of rCsHscB (5∼40 μg/ml), compared with medium or the production of *E. coli* transfected by empty vector (Fig. 5B, *P*<0.001). Furthermore, the production of IL-10 reached peak at the concentration of 20 μg/ml (almost 6 times increase). However, rCsHscB-induced IL-10 production in BMDM from TLR2 knockout mice was nearly abrogated (Fig. 5C).

**Figure 5.**
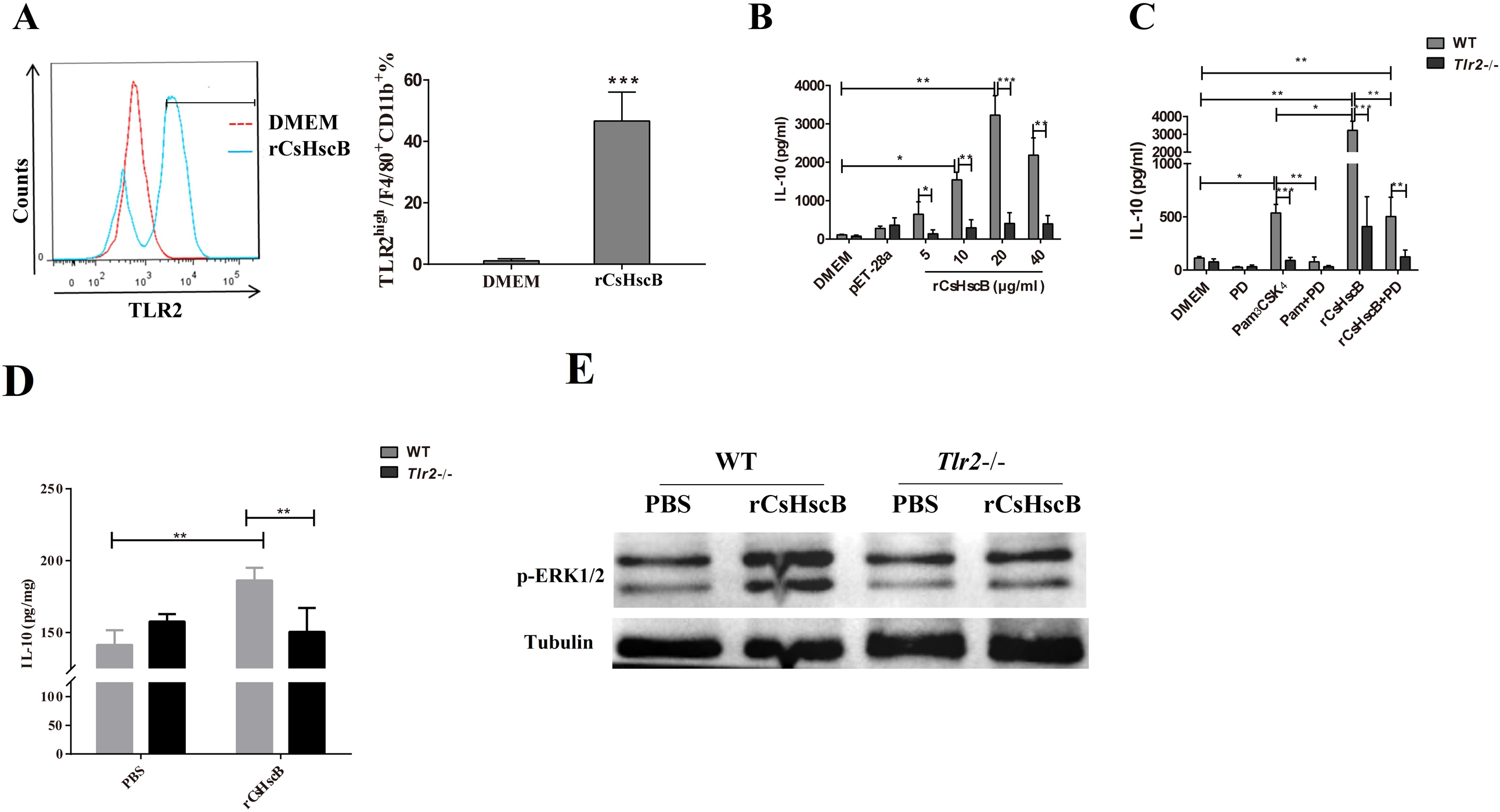
rCsHscB-induced IL-10 production depends on TLR2 *in vitro* and *in vivo*. **A** Flow cytometry analysis of TLR2 high F4/80^+^CD11b^+^ in DMEM stimulated and rCsHscB-stimulated BMDMs from wild type mice. **B** ELISA analysis of IL-10 production in BMDMs cells from wild type and *Tlr2*^-/-^ mice stimulated with various concentrations of rCsHscB as indicated. DMEM as negative control. pET-28 group represents supernatant of lysate from E. Coli transfecting with pET-28a empty vector. **C** ELISA analysis of IL-10 production in BMDMs cells from wild type and *Tlr2*^-/-^ mice stimulated with rCsHscB (20 μg/ml) with or without of ERK1/2 inhibitor (PD98059). PM3CSK4 were used as TLR2 ligand for positive controls. **D** The mice of wild type and *Tlr2* KO mice (∼20 g) were administrated with 20 μg rCsHscB or PBS for 24 h *i. v.*, and the mice were sacrificed and the liver from each mouse was collected for IL-10 detection. IL-10 production in supernatant of hepatic homogenate from each group was determined by ELISA. **E** Phosphorylation of ERK1/2 was attenuated in the liver of *Tlr2* KO mice that were received rCsHscB determined by western-blot. Quantitative data are representative of mean ± SEM of at least three independent experiments. Quantitative data are representative of mean ± SEM of at least three independent experiments. \**P* < 0.05, \*\**P* < 0.01, and \*\*\**P* < 0.001 compared with indicated group.

To verify whether rCsHscB-induced IL-10 production was depended on TLR2 mediated ERK1/2 signaling pathway, we used an inhibitor of ERK1/2 pretreated BMDM cells sourced from TLR2 wild type and TLR2 knockout mice and then stimulated by 20μg/ml rCsHscB for 24 h, IL-10 production in the culture were detected using ELISA. Again, the secretion of IL-10 in BMDM cells from TLR2 knockout mice was almost abolished when BMDM cells were stimulated by rCsHscB for 24 h (Fig. 5C). For TLR2 wild type BMDM cells, it showed that there was a significant decrease of IL-10 production in the BMDM cells with pretreatment of PD98059, compared with the cells pretreated by DMSO (the vehicle for PD98059). Furthermore, the data also demonstrated that the production of IL-10 was remarkably decreased (∼4 times decreased, Fig. 5C) in rCsHscB stimulated BMDM cells derived from *Tlr2*^-/-^ mice compared with that from *Tlr2* wild type mice. Collectively, our data demonstrated that rCsHscB induced IL-10 production in macrophage depends on the activation of TLR2-depended ERK1/2 signaling.

### rCsHscB could induce IL-10 in the liver of mice dependently by TLR2 mediated signaling pathway

To test whether rCsHscB could induce IL-10 production mediated by TLR2/ERK1/2 signaling pathway *in vivo* or not, the mice with or without *Tlr2* were both received rCsHscB (5 mg /kg body weight) or PBS by *i. v.* for 24 h, the levels of IL-10 in the hepatic homogenate were determined. The data showed that rCsHscB induced a higher level of IL-10 in the liver of mice, compared with the PBS group (Fig. 6A, *P*<0.01). However, IL-10 production in the liver from *Tlr2*^-/-^ mice were significantly lower than those in wild type mice when they were both received with the same dose of rCsHscB (Fig. 5D, *P*<0.01), but there was no any statistic difference in IL-10 in supernatant of hepatic homogenate in rCsHscB *Tlr2*^-/-^ mice and those from PBS treated *Tlr2*^-/-^ mice (Fig. 5D, *P*>0.05), suggesting that rCsHscB also induced IL-10 production in a TLR2 dependent manner *in vivo*. Furthermore, we also found that the phosphorylation of ERK1/2 in livers of *Tlr2*^-/-^ mice was also attenuated, compared with *Tlr2* wild type mice following administration of the same dose of rCsHscB (Fig. 5E). Collectively, these data demonstrated that rCsHscB could induce IL-10 production mediated by TLR2/ERK1/2 signaling pathway *in vivo*.

*C. sinensis* has evolved complex mechanisms for resistance to immune responses. Zhao et al demonstrated that total protein from *C. sinensis* inhibited Th1 immune responses by activation of mannose receptor (MR), but not TLR2 or TLR4 to induce Th2-skewed response [14]. Our previous study showed that TLR4 plays a regulatory role in the secretion of *C. sinensis* ESPs induced type I-relative cytokines (such as IFN-γ, IL-12, IL-6, TNF-α) [13]. However, the evidence suggests that the complex mechanisms for host-parasites interaction during *C. sinensis* infection are still poorly understood.

Many lipoproteins or lipo-peptide have been reported to display TLR2 ligands or agonists activities such as *Mycobacterium tuberculosis* (Mtb) LprG [5], Mtb LprA [23], *schistosomal* lyso-PS [8] and filarial Diacyl WoLP [9]. Thus, to identify the potential TLR2 agonist soured from *C. sinensis*, we screened the *C. sinensis* proteome data and predicted the potential lipoproteins using bioinformatic analysis. A lipoprotein from the family Co-chaperone Hsc20 (CsHscB) was ultimately selected for further study. However, it is very difficult to purify CsHscB directly from the worms due to lack of sufficient the background information as well as the low yield. We therefore used a recombinant CsHscB by *E. coli*, which was also recognized by sera of *C. sinensis* infect-mice, suggesting that rCsHscB remains the immunogenicity of *C. sinensis* rCsHscB. It was found that rCsHscB with the concentration of 5∼20 µg/ml could induce a strong production of IL-10 by macrophage in a dose-dependent manner. Similarly, it have been also demonstrated that recombinant PPE18 from M. tuberculosis or Pam_3_CSK_4_ known as the TLR2 ligands also trigger the activation of macrophages and production of IL-10 in a dosed manner by specifical interaction with TLR2 [24, 25]. Therefore, 20 µg/ml of rCsHscB was used as the optimized concentration for further study.

Pull-down assay is a useful approach to verify the protein-protein interaction *in vitro.* Using this assay, Chen et al. demonstrated that recombinant MPT83 derived from *M. tuberculosis* interacts specifically with TLR2 to promote the function of macrophage [26]. Our data suggested that rCsHscB soured from *C. sinensis* might acting as a TLR2 agonist plays a regulatory role in the immune responses to *C. sinensis* infection. However, the mechanisms by which TLR2 interact with rCsHscB are not known due to its complexity and further studies should be warranted.

During chronic infection, parasite products trend to induce strong regulatory responses which may be in charge of balanced host-parasite interaction whereby the tissues damages were impeded and worms’ survival was favored. IL-10 has been known as one of the mechanisms that contribute to induced regulatory responses induced by helminth infection [27]. For example, the increase production of IL-10 is mainly responsible for induction of CD4^+^ T cell hypo-responsiveness in the skin-draining lymph nodes after repeated exposure to *Schistosoma mansoni* larvae [28]. It is also evident that IL-10 sourced from CD4^+^CD25^−^ effector T cells impairs IFN-γ production for the control of acute inflammation and myositis in the diaphragm caused by *Trichinella spiralis* as well[29, 30]. In respect of *C. sinensis*, it showed that augment IL-10 was triggered by dendritic cells treated by *C. sinensis* crude antigen [14, 31]. Furthermore, it found that IL-10 secreted by lymphocytes from FVB was significantly higher than by those of BALB/c mice, which suggests that IL-10 may contribute to the susceptibility of different strains mice [21, 32]. In our present study, rCsHscB interacted with TLR2 can potently IL-10 production in macrophage with various concentrations (5∼20 μg/ml), which may account for mechanisms underlying production of IL-10 driven by *C. sinensis* infection.

Macrophage is one of the important sources of IL-10 in responses to TLRs or other PRRs ligands. But the intrinsic mechanisms that tailored production of IL-10 are still poorly understood. MAPKs signaling pathway (such as ERK, p38 etc) have been suggested to be involved in the control of the production of IL-10[33]. In our present study, we found that phosphorylated ERK but not p38 was activated in the responses to the stimulation of rCsHscB for 20∼30 min. Furthermore, the phosphorylated ERK1/2 was inhibited when the cells were pretreated by TLR2 blocking antibodies whereby the production of IL-10 was almost impaired, which suggest that the activation of ERK induced by rCsHscB in macrophage is mediated by TLR2. Interesting, although the production of IL-10 in rCsHscB induced cells was significantly decreased when the cells were pretreated by PD98059, these amounts were still higher than DMEM controls. To confirm these data, we also used another macrophage model-bone marrow-derived macrophage isolated from TLR2 wild and TLR2 knockout mice. Similar to our previous observation, the BMDM from TLR2 knockout mice impaired the secretion of IL-10 using the same amount of rCsHscB for stimulation, compared with those from *Tlr2*^-/-^ mice, but IL-10 secreted by rCsHscB-induced macrophages from TLR2 wild type mice were partial depressed when ERK1/2 inhibitor were pretreated. These data suggested that, in addition to ERK, other signaling pathways may be also involved in TLR2-dependent the production of IL-10 in macrophage-induced by rCsHscB. In addition, we also found that rCsHscB could induced the production of IL-10 via TLR2/ERK signaling pathway in mice following intraperitoneal injection with rCsHscB 5 mg/kg body weigh for 24 h, although there was limitation that the concentration of rCsHscB used the study might be not associated with real *C. sinensis* infection. To exclude any potential other effects of rCsHscB, we employed bull serum albumin (BSA) as a control for *in vivo* study, the data showed that BSA had little effects on the production of IL-10 as well as the activation of p-ERK *in vivo* (the data is not shown), which demonstrated rCsHscB could specially induce high level of IL-10 both *in vivo* and *in vitro.*

In conclusion, in the present study, we identified CsHscB sourced from *C. sinensis* acting as a novel TLR2 agonist to induce potently activation of macrophage, our study also demonstrates that a robust IL-10 production by rCsHscB-induced macrophage is dependent by TLR2-mediated ERK1/2 signaling pathway, which may reveal a novel mechanism for host-parasites recognition during *C. sinensis* infection. The present study will contribute to a better understanding of the interaction between the worms and host cells. In addition, rCsHscB might be suggested in the development of novel therapeutic strategies with implications beyond parasitology due to its potential regulatory capacities of immune responses.

## Author Contributions

CY and KYZ conceived and designed the experiments. FF and ZYZ performed the majority of experiments. YZ, JW, DX, HLL, YGW, QY and RXT contributed to the acquisition of data. CY and ZYZ wrote the paper. All authors read and approved the final manuscript.

